# Extending Levelt’s Propositions to perceptual multistability involving interocular grouping

**DOI:** 10.1101/057232

**Authors:** Alain Jacot-Guillarmod, Yunjiao Wang, Claudia Pedroza, Haluk Ogmen, Zachary Kilpatrick, Krešimir Josić

**Affiliations:** Dept. of Electrical&Computer Engineering, University of Houston, Houston, Texas, 77204 USA; Dept. of Mathematics, Texas Southern University, Houston, Texas, 77204 USA; Center for Clinical Research and Evidence-Based Medicine, McGovern Medical School at The University of Texas Health Science Center at Houston, Houston, Texas, 77204 USA; Dept. of Mathematics, University of Houston, Houston, Texas, 77204 USA; Dept. of Applied Mathematics, University of Colorado, Boulder, Colorado, 80309, USA; Dept. of Biology and Biochemistry, University of Houston, Houston, Texas, 77204, USA; Dept. of Bioscience, Rice Univer—sity, Houston, Texas, 77251 USA; Lausanne University, Hospital Lausanne, Switzerland

**Author notes:** Jacot-Guillarmod and Wang are co-first authors, and Josić and Kilpatrick are co-corresponding authors.

**Keywords:** Multistable perceptual rivalry, Levelt’s propostions, interocular grouping

## Abstract

Levelt’s Propositions have been a touchstone for experimental and modeling studies of perceptual multistability. We asked whether Levelt’s Propositions extend to perceptual multistability involving interocular grouping. To address this question we used split-grating stimuli withcomplementary halves of the same color. As in previous studies, subjects reported four percepts in alternation: the two stimuli presented to each eye (single-eye percepts), as well as two interocularly grouped, single color percepts (grouped percepts). Most subjects responded to increased color saturation by more frequently reporting a single color image, thus increasing the predominance of grouped percepts (Levelt’s Proposition I). In these subjects increased predominance was due to a decrease in the average dominance duration of single-eye percepts, while that of grouped percepts remained largely unaffected. This is in accordance with generalized Levelt’s Proposition II which posits that the average dominance duration of the stronger (in this case single-eye) percept is primarily affectedbychanges in stimulus strength. In accordance with Proposition III, the alternation rate increased as the difference in the strength of the percepts decreased. To explain the mechanism behind these observations, we introduce a hierarchical model consisting of low-level neural populations, each responding to input at a visual hemifield, and higher-level populations representing the percepts. The model exhibits the changes in dominance duration observed in the data, and conforms to all of Levelt’s Propositions.

## 1. Introduction

The brain is remarkably adept at interpreting noisy and ambiguous visual inputs (Kersten et al., 2004; Fiser et al., 2010). However, sometimes competing interpretations of a stimulus are not disambiguated, and different interpretations are perceived in alternation. For example, binocular rivalry occurs when the two eyes are presented with disparate images. Instead of perceiving a fusion of the two images, one experiences intermittent switching between two distinct percepts (Wheatstone, 1838; Blake and Logothetis, 2002). Multistable perceptual phenomena have been used extensively to study visual awareness and its underlying cortical mechanisms (Leopold and Logothetis, 1996; Polonsky et al., 2000; Tong et al., 2006; Sterzer et al., 2009).

**Figure 1:**
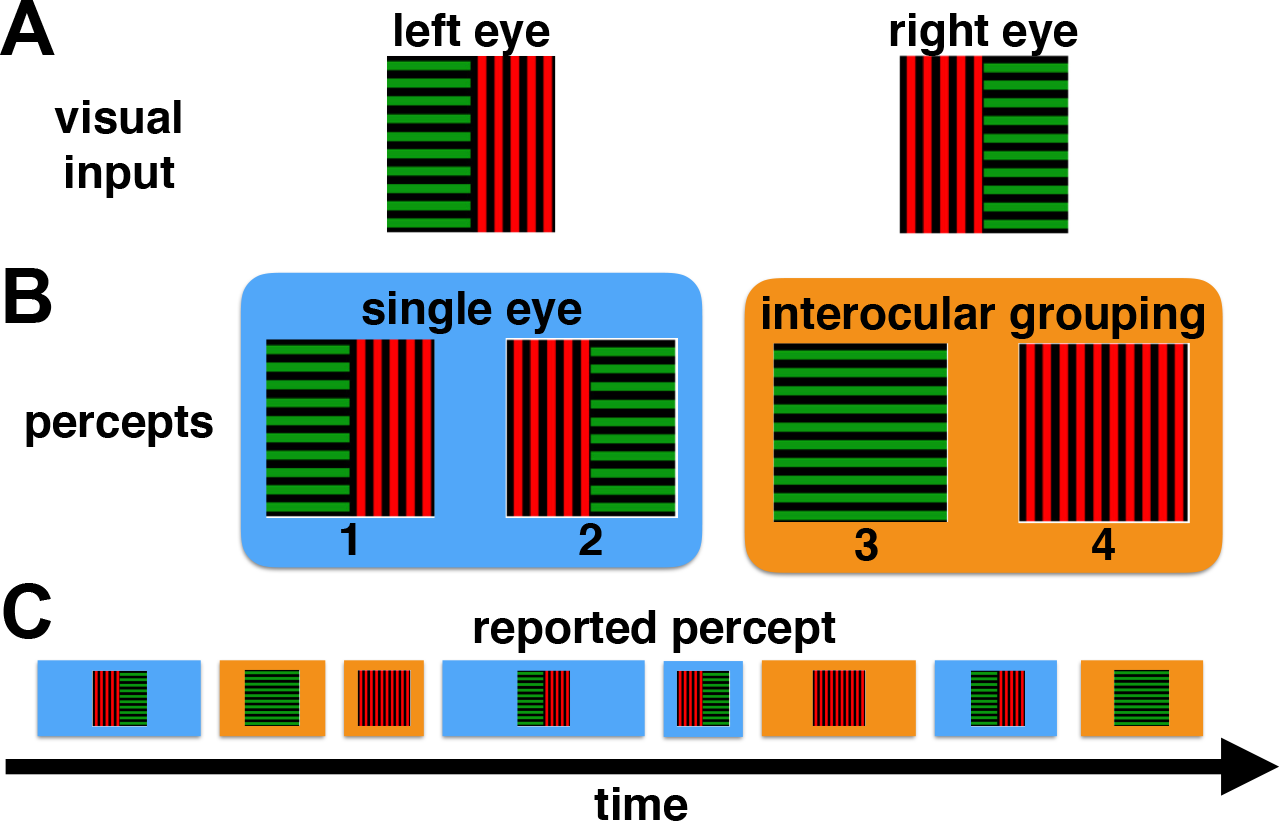
(A) An example of the stimuli presented to the left and right eyes. Gratings were always split so that halves with the same color and orientation could be matched via interocular grouping, but were otherwise randomized across trials and blocks (see Methods). (B) Subjects typically reported seeing one of four percepts-two single-eye and two grouped-at any given time during a trial. (C) A typical perceptual time series reported bya subject, showing the stochasticity in both the dominance times and the order of transitions between percepts.

Levelt’s observations (Levelt, 1965) have become a touchstone for experimental and modeling studiesof perceptual rivalry (Blake, 1989; Moreno-Bote et al., 2007; Shpiro et al., 2007; Wilson, 2003; Said and Heeger, 2013; Seely and Chow, 2011). Levelt’s original Propositions relate *stimulus strength, predominance* (the fraction of time apercept is dominant), and *dominance durations* (the duration of the dominant percept) in binocular rivalry (Brascamp et al., 2015). However, generalizations of these propositions have been shown to hold in many cases of perceptual multistability (Bossink et al., 1993; Bonneh et al., 2001; Brascamp et al., 2006; Moreno-Bote et al., 2010; Klink et al., 2008).

We hypothesized that Levelt’s propositions also extend to perceptual multistability in the case of interocular grouping (Kovacs et al., 1996) where observers report alternation between percepts that combine features of two disparate images presented simultaneously to the two eyes (Tong et al., 2006; Diaz-Caneja, 1928). For instance, when multiple patches of two visual scenes are intermingled and the results presented to different eyes, observers intermittently perceive the original, coherent scenes (Kovacs et al., 1996). In this case conflicting percepts arise via interocular grouping-the binding of portions of the visual input from both eyes into acoherent percept.

Levelt’s Propositions generalize naturally to this case: (I) Increasing percept strength of grouped percepts increases the perceptual predominance of those percepts. (II) Increasing the difference between the percept strength of grouped percepts and that of single-eye percepts increases the average perceptual dominance duration of the stronger percepts. (III) Increasing the difference in percept strengths between grouped percepts and single-eye percepts reduces the perceptual alternation rate. (IV) Increasing percept strength in both grouped percepts and single-eye percepts while keeping it equal among percepts increases the perceptual alternation rate Brascamp et al. (2015). Here we refer to *percept* strength rather than *stimulus* strength since both single-eye and grouped percepts share the same visual inputs.

To test our hypothesis we used split-grating stimuli (See Fig. 1A) for which subjects reliably reported four percepts in alternation: single-eye percepts-the two stimuli presented to each eye (percepts 1 and 2 in Fig. 1B), as well as two interocularly grouped, single color, coherent percepts (3 and 4 in Fig. 1B). We hypothesized that an increase in color saturation increases the strength of the coherent, grouped percepts. Indeed, we found that for most subjects an increase in color saturation lead to increased predominance of grouped percepts (Proposition I). At the same time the dominance duration of single-eye (stronger) percepts decreased, while that of grouped (weaker) percepts remained largely unaffected (Proposition II). As a consequence, the alternation rate increased with a reduction in the difference of percept strengths (Proposition III). In addition, we found that an increase in the predominance of grouped percepts was partly due to an increase in the fraction of visits to grouped percepts.

We next investigated whether classical models of binocular rivalry can explain the neural mechanisms behind the present form of perceptual multistability. To do so we developed a hierarchical model consisting of four low-level populations responding to input in each hemi-field of the two eyes, and higher-level populations representing the four percepts (Wilson, 2003; Diekman et al., 2013). We assumed that an increase in color saturationincreased interocular coupling strength between low-level neural populations responding to complementary stimuli. Our model showed that the same neural mechanisms (*i.e.* mutual inhibition, adaptation and noise) used to explain binocular rivalry Laing and Chow (2002) also explain the main features of our experimental findings. In addition, the behavior of the model conformed with Proposition IV, which we were not able to test experimentally.

Interestingly, the level of change in color saturation in our experiments did not result in changes in interocular grouping in all subjects. However, when the effect was present, it could be explained by the generalization of Levelt’s propositions.

## 2. Methods

### 2.1. Experiment

*Observers.* Nine observers with normal or corrected-to-normal vision, including three of the authors (AJ, ZK, YW), participated in this experiment. Six were naive to the experimental hypotheses and three were not. The experiments were conducted according to a protocol approved by the University of Houston Committee for the Protection of Human Subjects and in accordance with the federal regulations 45 CFR 46, the ethical principles established by the Belmont Report, and the principles expressed in the Declaration of Helsinki. All participants provided their written informed voluntary consent following the consent procedure approved by the University of Houston Committee for the Protection of Human Subjects. Data are presented for all nine subjects.

*Apparatus.* The visual stimuli used in the experiment were generated using a VSG visual stimulus generator card (VSG 2/5, Cambridge Research Systems). The stimuli were displayed on a calibrated 19” high resolution color monitor with a 100 Hz frame rate. Monitor calibration was carried out using CRS colorCAL colorimeter. A head/chin rest was used to stabilize observers’ head position. The distance between the monitor and the observer was set to 108 cm. We used a stereoscopic mirror arrangement (haploscope) in order to present the left and right stimuli separately to the left and right eyes. It consisted of four mirrors, whose horizontal/vertical positions and inclinations could be adjusted using screws.

*Stimuli.* Subjects were presented with variations of the stimulus depicted in Fig. 1A. A square composed of two orthogonal gratings was presented to each eye using the haploscope. The orthogonal gratings were arranged so that interocular grouping resulted in a percept with single, i.e., uniform orientation (horizontal or vertical). In order to have a stimulus parameter to control the percept strength for this interocular grouping, we have added color to our stimuli, such that interocular grouping would lead not only to a uniform orientation but also to a uniform color (Fig. 1A). Stalmeier and de Weert (1998) studied the contribution of color and luminance contrast to binocular rivalry. In their experiments, the stimulus to one eye was achromatic concentric rings whereas the stimulus to the other eye was a radial pattern made of isoluminant color pairs. They showed that the dominance duration of the colored radial pattern, hence the strength of the chromatic input, increased as the chromatic distance, *d(u,v)*, between the colors in the CIE 1960 space increased up to *d(u,v)*≈0.1, and saturated thereafter. There were also significant differences in dominance durations depending on the criterion for isoluminance (flicker photometry vs minimal distinct border (MDB) criterion), and the direction of change in the color space. Finally, their results showed inter-subject variability both in the effectiveness of pure chromatic contrast and achromatic contrast.

In preliminary observations, we found color saturation effectively controlled percept strength for interocular grouping. Hence, grating halves were assigned a color-either red or green-at two different saturation levels, 0.4 or 0.9. The HSV color space coordinates for red and green were (0.497, 0.4/0.9, 0.7) and (120.23, 0.4/0.9, 0.7), respectively, with the pair of values 0.4/0.9 referring to two different levels of color saturation. Atlow saturation (S=0.4), the corresponding CIE 1960 (u,v) coordinates for red were (0.214, 0.3) and L=57.7*cd/m*^2^; whereas for green they were (0.169, 0.315) and L=72*cd/m*^2^. At high saturation (S=0.9), the corresponding CIE 1960 (*u,v*) coordinates for red were (0.333, 0.329) and L=25.4*cd/m*^2^ whereas for green they were (0.127, 0.360) and L=57.6*cd/m*^2^. At low saturation, the chromatic distance *d(u, v)* between the two colors was *d(u, v)*=0.05 and the achromatic distance in terms of Michelson Contrast (MC) was MC=0.11. At high saturation, these values were *d(u,v)*=0.21 and MC=0.388. Hence, by changing color saturation from 0.4 to 0.9, stimulus strength was increased significantly both in chromatic and achromatic dimensions. It is also noteworthy that the chromatic distance values of 0.05 and 0.21 fall to the left and right of the critical distance *d(u,v)*≈0.1 at which the strength of the chromatic stimulus for binocular rivalry starts to saturate as observed by Stalmeier and de Weert (1998).

To allow for interocular grouping of complementary patches, the two halves with the same orientation always shared the same color at the same saturation level, and were shown to opposite hemifields of either eye. For example, the combination horizontal green/vertical red presented to the lefteye determined the combination vertical red/horizontal green presented to the right eye, as well as the two grouped percepts-vertical red and horizontal green (See Fig. 1B). In total,there were four possible stimulus arrangements, all completely determined by any half of a stimulus presented toone eye. The two squares were displayed on a grey background (0.0, 0.0, 0.2): *(u,v)*=(0.188, 0.442) and L=23.88*cd/m*^2^ and were contained within a square frame with a protruding horizontal and vertical line to help image alignment.

*Experimental procedure.* Each *session* was divided into six 3-minute trials separated by a 90-second resting period. To account for the time it took subjects to adjust to the stimuli and form stable percepts, the first 30 seconds of each trial were not analyzed. The association between color and orientation was maintained within a single session, but was randomized across sessions. For example, we used a vertical red/horizontal green left eye stimulus across some sessions (Fig. 1A). In contrast, saturation and the position of the horizontal grating was randomized across the six trials. Within one session, each saturation level appeared in three trials and each grating positioning occurred in three trials.

Four subjects finished 6 total sessions (AJ, MA, ZK, ND), three subjects finished 5 sessions (FG, YW, ML), one subject finished 4 sessions (AB) and the remaining one finished 7 sessions (ZM). Therefore, after discarding the initial 30 seconds of each trial, a total of about 90 minutes of data over about 36 trials was collected per subject: about 18 trials for each saturation conditions, with 3 trials per level and color/orientation pairing. See the Supplementary Material which has been deposited to Github (https://github.com/YunjiaoWang/multistableRivalry.git) for more details. Subjects were asked to indicate the dominant percept by holding down one of four different buttons (1, 2, 3, 4) on a gamepad. They were instructed to press button 1 when perceiving a split grating with left part red; button 2 when perceiving split grating with left part green; button 3 when perceiving an all red grating; and button 4 when perceiving an all green grating. When the perceived image did not correspond to one of these four options, subjects were instructed to release all buttons. Such a report typically marked a transition between percepts, but could also be followed by a transition to the same percept. Before the beginning of the experiment, subjects were familiarized with the controller.

### 2.2 Data analysis

We performed the statistical analysis in R and provide a description ofthe analysis below. Commented code, as well as all collected data are available in the Supplementary Material.

We conducted all data analyses under a Bayesian framework. Standard significance tests would allow us to reject the null hypothesis that a color saturation change has no effect on dominance time, but would not allow us to accept the alternative hypothesis. In contrast, a Bayesian approach allows us to conclude that for some subjects a change in color saturation did affect percept dominance. We believe that showing the probabilities that this effect was present is more informative than concluding that a null hypothesis is rejected at some (arbitrary) significance level. Our use of Bayesian statistics means that confidence intervals are replaced by credible intervals, and traditional notions of “significance” do not apply. Instead of using a fixed threshold for significance, we provide the probabilities that a change in color saturation affects the perception ofthe stimuli, given the data (Wasserstein and Lazar, 2016).

Importantly, in our analysis we use a hierarchical model to analyze concurrently the data from all subjects in the experiment (Gelman and Hill, 2006). Such models address the issue of multiple comparisons and provide efficient estimates (Gelman et al., 2012).

*Predominance of grouped percepts.* Using the time series recorded from each trial, we computed the predominance of grouped percepts. Predominance is the fraction of time that subjects reported a grouped percept, Tg_rouped_, by pressing the corresponding gamepad button, out of the total time they reported any percept (percepts 1, 2, 3 or 4), *i.e.*

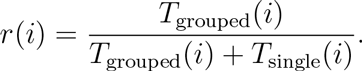

Here i is the number of the trial, with 18 trials at each color saturation level (0.4 and 0.9). This is equivalent to the fraction of time that buttons 3 or 4 were pressed out of the total time any button was pressed during trial i. In our analysis, we partitioned trials based on the color saturation level used for each trial, grouping across all other conditions. We analyzed changes in predominance using a linear Student-t regression model to account for skewness in the data. We included the condition (low/high color saturation) as a covariate and set the degrees of freedom of the t distribution to 4 to provide robust inference while avoiding computational difficulties often encountered when using a prior for the degrees of freedom (Fonseca et al., 2008). Letting *r_ij_* be the predominance for subject *j* in trial *i*, the model is specified as:

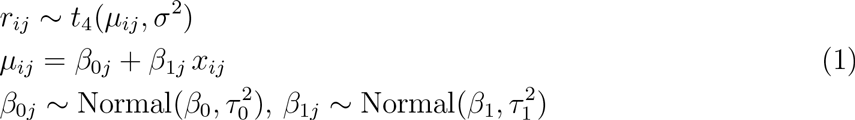
where *x_ij_* is the color saturation indicator (1 for 0.9, 0 for 0.4). The random regression coefficients β_0j_ and β_1j_allow the effects of color saturation to vary across subjects. This hierarchical model assumes that the effects from different subjects are similar but not identical and come from the same population with overall means of β_0_ and β_1_. Prior distributions for the overall saturation effects 0_O_ and were independent and normal with mean 0, and variance 10^4^. We used Uniform(0, 100) priors for the standard deviation of the random effects, *T*_0_ and *t*_1_ and Uniform(0, 1000) for σ. We estimated the mean difference in the fraction of time between the two saturation levels and its 95% credible interval (CI) and the probability that the difference is greater than 0. We performed an equivalent analysis to examine whether the mean dominance time of the single eye or grouped percepts changed across conditions.

From the *i^th^* trial in each condition, we also computed ratios of the number of visits to grouped percepts, *N*_grouped_, over the number of all visits to either single-eye or grouped percepts,

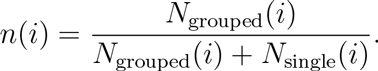

We used the model specified in Eq. (1) to analyze *n(i)* and determine the change in the fraction of visits to the grouped or single-eye percepts across conditions.

*Transition probabilities.* To estimate the transition probabilities between percept types, we classified percepts into two states: single-eye, *S*, corresponding to percepts 1 and 2, and grouped, *G*, corresponding to percepts 3 and 4. For each trial, we converted the data into two binary sequences: One sequence contained all transitions from state S with transitions from *S* to *S* denoted by 1, and from *S* to *G* by 0. The second sequence contained transitions from *G*, those from *G* to *G* denoted by 1, and from *G* to *S* by 0. We used all data obtained by each subject in a given condition (low/high color saturation)to estimate the transition probability from *S* to *S*, and from *G* to *G*. The model isspecified as

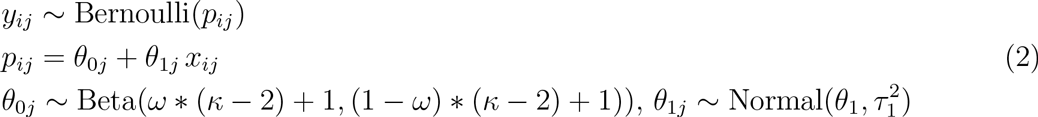
where *x_ij_* is the color saturation indicator (1 for 0.9, 0 for 0.4). We used vague priors: a uniform prior on the interval [0,1] for the mode, ω, and a Gamma prior with rate and shape both equal to 0.01 for the concentration parameter, *κ*. Prior distributions for the overall saturation effects θ_1_ was independent of these, and normal with mean 0, and variance10^4^. We used Uniform(0, 100) prior for the standard deviation of the random effect *T*_1_.

*Model implementation.* All Bayesian models were implemented via Markov Chain Monte Carlo methods in JAGS. We used 3 MCMC chains with at least 20,000 iterations after an initial burn-in of 4000 iterations. We assessed convergence by calculating the Gelman-Rubin diagnostic, 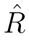 for all parameters.

### 2.3 Model formulation and simulation

To provide a mechanistic explanation of our observations we used a hierarchical neural population model, based on previous work (Laing and Chow, 2002; Wilson, 2003, 2009; Moreno-Bote et al., 2007; Huguet et al., 2014; Diekman et al., 2013). A schematic representation of the model is shown in Fig. 2. The sub-network at the first level of the hierarchy consists of four neural populations, each receiving input from a different hemifield of the two eyes (See also Fig.6C of Diekman et al. (2013) and Fig.2B of Tong et al. (2006)).

The responses of all four possible pairs of populations at the first level, corresponding to complementary hemifields, are integrated by distinct populations at the second level (Laing and Chow, 2002; Wilson, 2003; Moreno-Bote et al., 2007). Each of these four pairs corresponds to one of the four percepts shown in Fig. 1B, and thus each second level population can be associated with a distinct percept. We assumed *excitatory* coupling between populations receiving input from different hemifields both from the same and from different eyes. We also assumed *inhibitory* coupling between populations receiving input from the same hemifield of different eyes, *e.g.* the left hemifield of the left and the left hemifield of the right eye. This is consistent with electrophysiology and tracing experiments that reveal long-range horizontal connections between cells with non-overlapping receptive fields, and similar orientation preferences (Stettler et al., 2002; Sincich and Horton, 2005). Moreover, cells with orthogonal orientation preferences can inhibit one another through recurrent and feedback circuitry (Ringach et al., 1997; Ferster and Miller, 2000). Finally, we assumed that all populations at the second level inhibit each other. This is consistent with previous computational models which have recapitulated results of psychophysics experiments for rivalry between two percepts (Lankheet, 2006; Seely and Chow, 2011).

**Figure 2:**
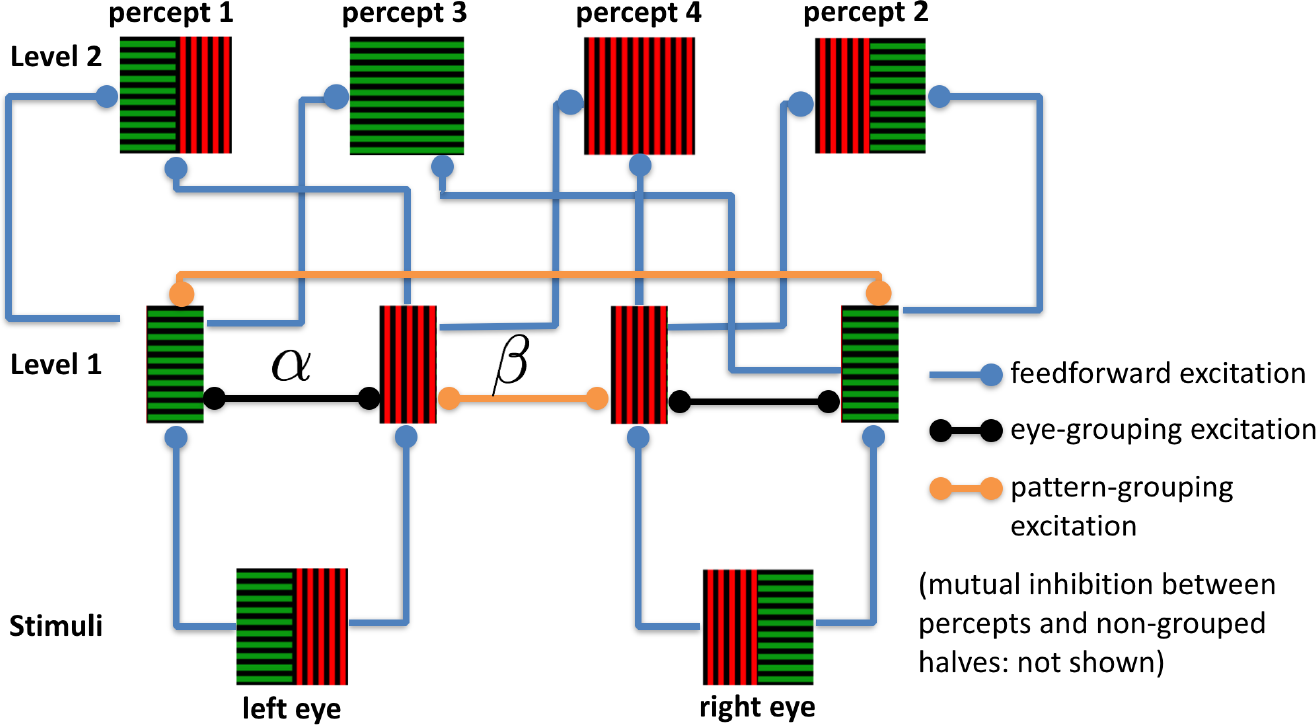
Computational model of interocular grouping. Neural populations representing stimuli to the four hemifield-eye combinations at Level 1 provide feedforward input to populations representing integrated percepts at Level 2, as described by Eqs. (3) and (5) (See also Fig. 6C of Diekman et al. (2013) and Fig. 2B of Tong et al. (2006)). Recurrent excitation within Level 1 is shown, whereas mutual inhibition between the same hemifield of opposite eyes is not shown. All populations in Level 2 mutually inhibit one another (Laing and Chow, 2002;Wilson, 2003; Moreno-Bote et al., 2007).

The two levels thus form a processing hierarchy (Wilson, 2003; Tong et al., 2006) with the first roughly associated with monocular neural activity generated in LGN and V1 (Wilson, 2003; Blake, 1989; Polonsky et al., 2000; Tong, 2001), and the second level associated with the activity of higher visual areas, such as V4 and MT, that process objects and patterns (Leopold and Logothetis, 1999; Wilson, 2003; Lamme and Roelfsema, 2000). However the activity dynamics described by each level is likely to correspond toa distributed process which spans multiple functional layers of the visualsystem (Sterzer et al., 2009).

*Equations describing Level 1.* The activity of each neural population receiving input from one of the four hemifield-eye combinations at Level 1 is described by a firing rate variable *E_i_*, *i*=1, 2, 3,4 (corresponding to left hemi/left eye; right hemi/left eye; left hemi/right eye; and right hemi/right eye, see Fig. 2). These rates are governed by the following equations:

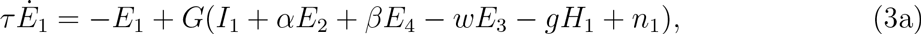

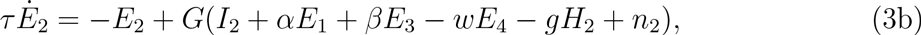

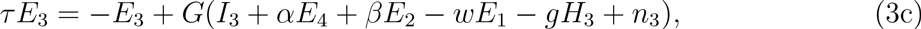

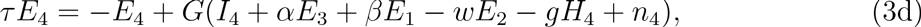
with the activity time constant *τ*=10ms (Häusser and Roth, 1997), *I_i_* is drive from the stimulus, *H_i_*is used to model rate adaptation with strength *g*, and *n_i_* models random fluctuations due to network effects and synaptic noise (Faisal et al., 2008). The strength of within eye excitatory coupling is determined by α, while cross-eye excitatory coupling between populations receiving input from complementary hemifields is described by β. The strength of mutual inhibition due to orientation and color competition is determined by *w*. We used a sigmoidal gain function, *G*, to relate the total input to the population to the output firing rate:

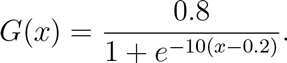

This choice was not essential, as we could have used other gain nonlinearities, such as a Heaviside step or a rectified square root, as long as each individual population, *E_i_*, possesses both a low and high firing rate state (Laing and Chow, 2002; Moreno-Bote et al., 2007).

The rate adaptation variables, *H_i_* describe the population-wide effects of hyperpolarizing currents, activated due to increases in firing rates (Benda and Herz, 2003),

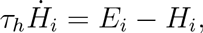
for *i*=1, 2, 3, 4 with time constant τ_h_=1000ms. Following Moreno-Bote et al. (2007), we modeled the fluctuations in population input currents with an Ornstein-Uhlenbeck process,

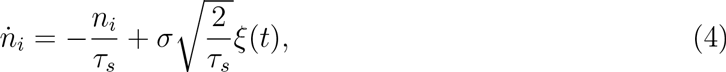
where *τ_s_*=200ms, σ=0.03, and ξ(t) is a white-noise process with zero mean. Changing the timescale and amplitude of noise does not impact the results significantly.

*Equations describing Level 2.* The activity of the neural populations associated with percept *i* at Level 2 is described by the mean firing rate *P_i_* for *i*=1, 2, 3, 4. The *P_i_* are governed by

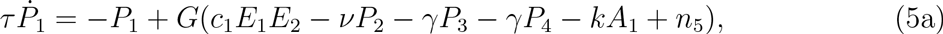

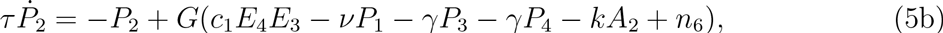

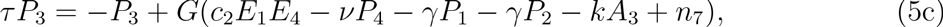

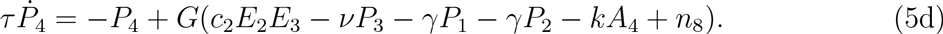

We model the feedforward inputs to each second level population as the product of the activities, *E_j_*,*E_k_*, of the associated populations at the first level. For instance, activity *P_1_* depends on the product *E*_i_*E*_2_ since percept 1 iscomposed of the stimuli in the hemifields providing input to populations 1and 2 at the first level. Previous experimental and mod-eling studies havepointed to such multiplicative combinations of visual field segments as a potential mechanism for shape selectivity (Salinas and Abbott, 1996; Brincat and Connor, 2006). Again, we model rate adaptation using a separate variable, *A_i_*, described by

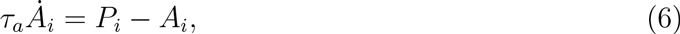
where we set *T_a_*=*T_h_*. When we replaced the multiplicative input to the second level population with additive input from Level 1, *E_j_*+*E_k_*, our results remainedqualitatively unchanged.

Theorem 2.2 in Diekman et al. (2012) suggests that when β=0 and α is large enough, the lower level network model can be reduced to a classical mutual inhibitory two-node model. Here, we choose the parameter ranges of α and *I* so that all four Levelt’s propositions hold when β=0.

**Figure 3:**
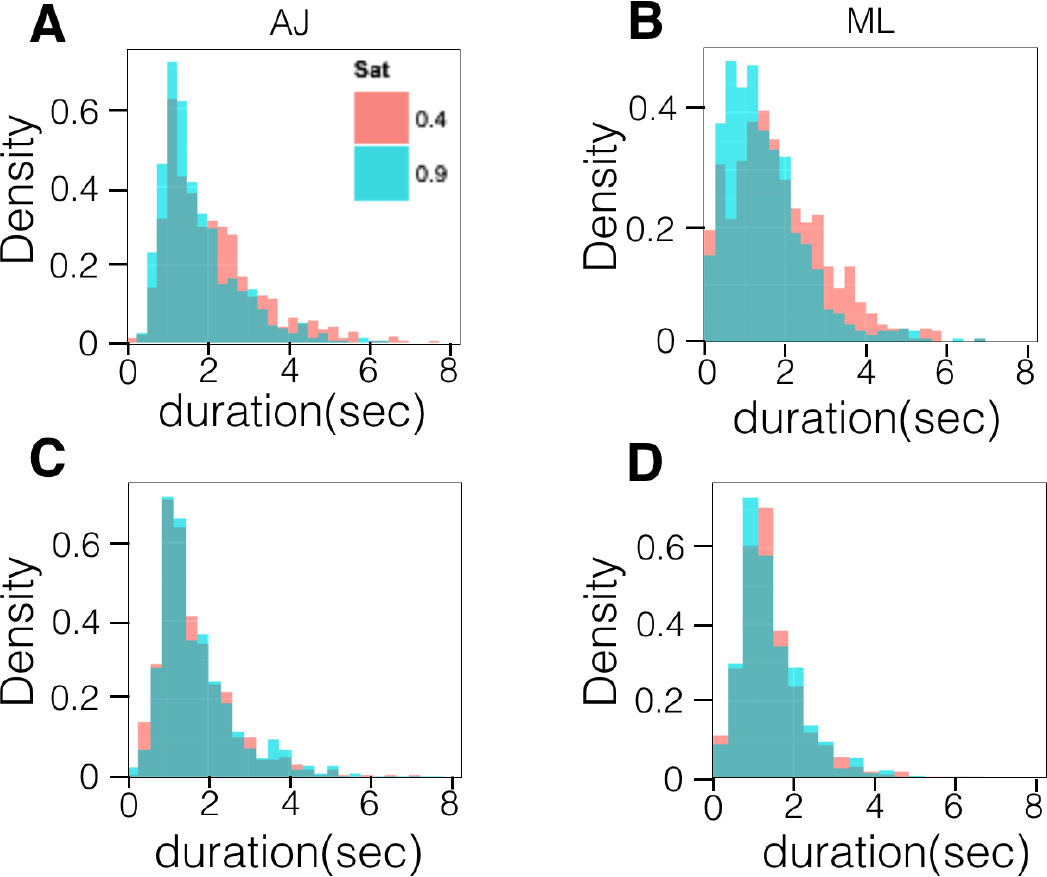
Dominance times for two subjects, ML and AJ, approximately follow a gamma distribution. (A,B) Histograms of single-eye percept durations are unimodal, but somewhat different between the two saturation conditions. (C,D) Histograms of the grouped percept durations are closer to each other. Each histogram contains data collected from 18 trials of 2.5 minutes each, amounting to approximately 1200 dominance duration reports (See Methods and Supplementary Material for more details).

## 3. Results

Nine observers were presented with two split-grating images simultaneously to each eye using a haploscope (See Methods). Subjects reported one of four possible percepts by pressing buttons on a game pad. We examined how the fraction of time subjects perceived grouped images (the *predominance* of grouped images) depended on the color saturation of the stimuli. We observed an effect in some subjects, and propose a model to explain it.

*Dominance durations follow a gamma distribution.* We computed dominance duration as the total time that a subject reported seeinga percept, *i.e.* the total time that the subject continuously pressed a button on the gamepad, corresponding to a percept. For all subjects the distribution of dominance times for single-eye and grouped percepts had the shape of a gamma distribution. This is consistent with previous studies of perceptual multistability (Blake and Logothetis, 2002; Brascamp et al., 2005; van Ee, 2009). For some, but not all subjects, the mean of single-eye percept times decreased with an increase in color saturation (Fig. 3). A more thorough analysis was therefore needed to determine the effect of color saturation on percept predominance.

*Predominance of grouped percepts.* We first examined whether an increase in color saturation affected the fraction of time grouped percepts were reported. Our hypothesis was that predominance of grouped percepts increases with color saturation, as a result of a stronger visual cue to bind the two complementary halves of the stimuli presented to each eye into a coherent percept (Wagemans et al., 2012). The data supports this effect in five out of nine subjects (Fig. 4): A Bayesian analysis of the data shows that for five out of the nine subjects there was a 0.92 or higher probability that the difference in mean predominance times increased between the conditions, given the reported percept durations (See Table in Fig. 4). There was no evidence that changes in color saturation impacted predominance in the remaining subjects. We next examined how this change in predominance was related to both changes in average dominance time and the frequency of visits to single-eye versus grouped percepts.

**Figure 4:**
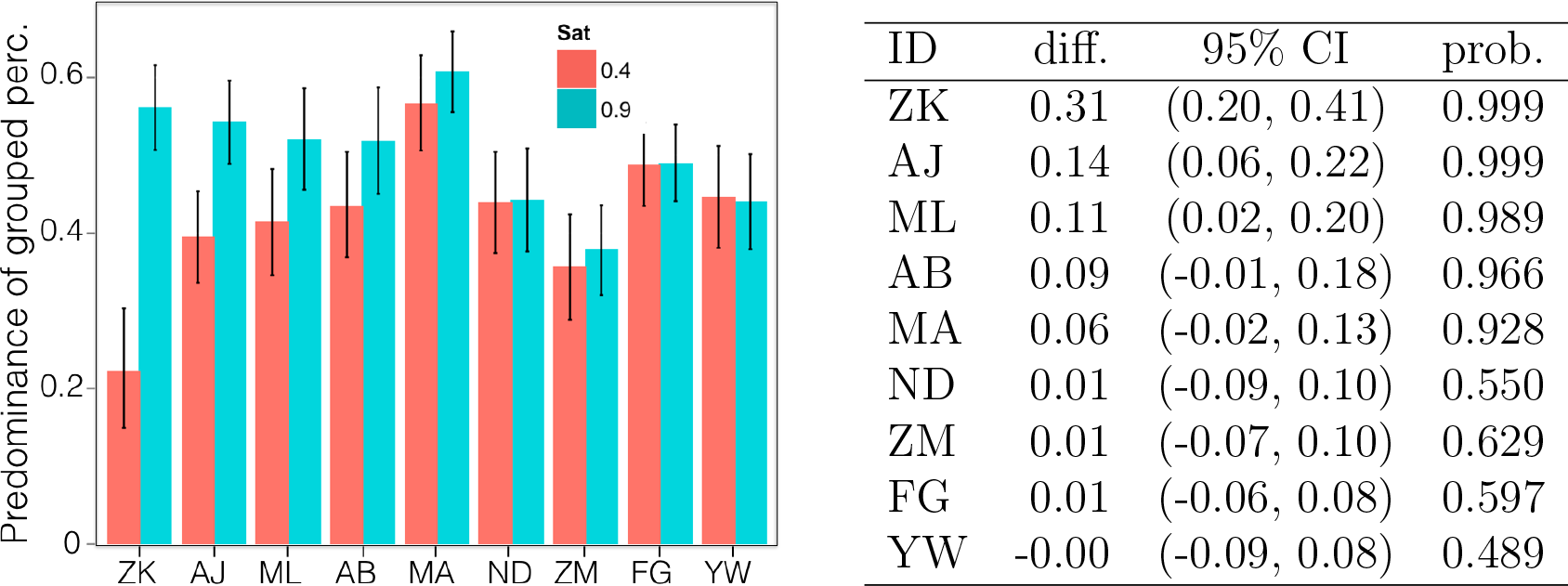
(Plot) Grouped percept predominance: each colored bar indicates the mean predominance at a given color saturation level in a given subject and black error bars denote the 95% credible intervals. (Table) Differences between ratios at the two color saturation levels: diff.=difference of predominance means at saturation 0.9 and 0.4; 95% CI stands for 95% credible interval; ‘prob.’ is the probability that the predominance of grouped percepts is higher at saturation level 0.9 (See Methods). We use the same ordering of subjects in all subsequent tables and figures, so that the five subjects sensitive to changes in color saturation are listed first.

## 3.1. Causes of predominance changes

In the case of only two percepts, the number of visits to each percept will differ by at most one per trial (van Ee, 2009), and dominance duration is closely related to predominance. With more than two percepts, they donot simply alternate. The order in which multiple percepts appear affects predominance (Naber et al., 2010; Huguet et al., 2014). Thus, to understand changes in predominance we must examine how color saturation influences dominance duration, as well as the number of visits to each percept.

*Single-eye percept durations decrease with color saturation.* We compared the average dominance durations of single-eye and grouped percepts for the two different color saturation conditions in Fig. 5. In six out of nine subjects, there was a higher than 0.95 probability that dominance duration of single-eye percepts decreased as color saturation increased (subjects ZK, AJ, ML, AB, MA, ZM, See Fig. 5A). These included the five subjects for which the predominance of grouped percepts increased. There was no strong evidence that increased color saturation increased the dominance duration of grouped percept in any subjects. The distribution of dominance durations for two subjects in Fig. 3 show a decrease in the mean dominance duration of single-eye percepts (panels A and B) with little change in grouped percept durations (panels C and D).

Generalized Levelt’s Proposition II states that increasing the difference between the percept strength of grouped percepts and that of single-eye percepts primarily increases the average perceptual dominance duration of the stronger percepts Brascamp et al. (2015). By increasing color saturation, we decreased the difference in stimulus strength between single-eye and grouped percepts: In the low color saturation case, the single-eye percepts were stronger, as their predominance was higher than that of grouped percepts (Fig. 4, for seven of the nine subjects the predominance of grouped percepts was below 0.5 with a probability of 0.94 or higher. See Supplementary Material). At higher color saturation the grouped percepts had a mean predominance of near 0.5 or below for eight subjects. We also note that grouping during binocular rivalry is dictated by the eye-of-origin (Stuit et al., 2014), so it is reasonable to assume that single-eye percepts remain stronger even at higher color saturation. Thus, for most subjects who were sensitive to a change in percept strength the stronger percepts’ (single-eye) mean dominance duration decreased, while the weaker percepts’ (grouped) durations remained roughly the same. We explore further comparisons with Propositions II-IV in the Discussion.

**Figure 5:**
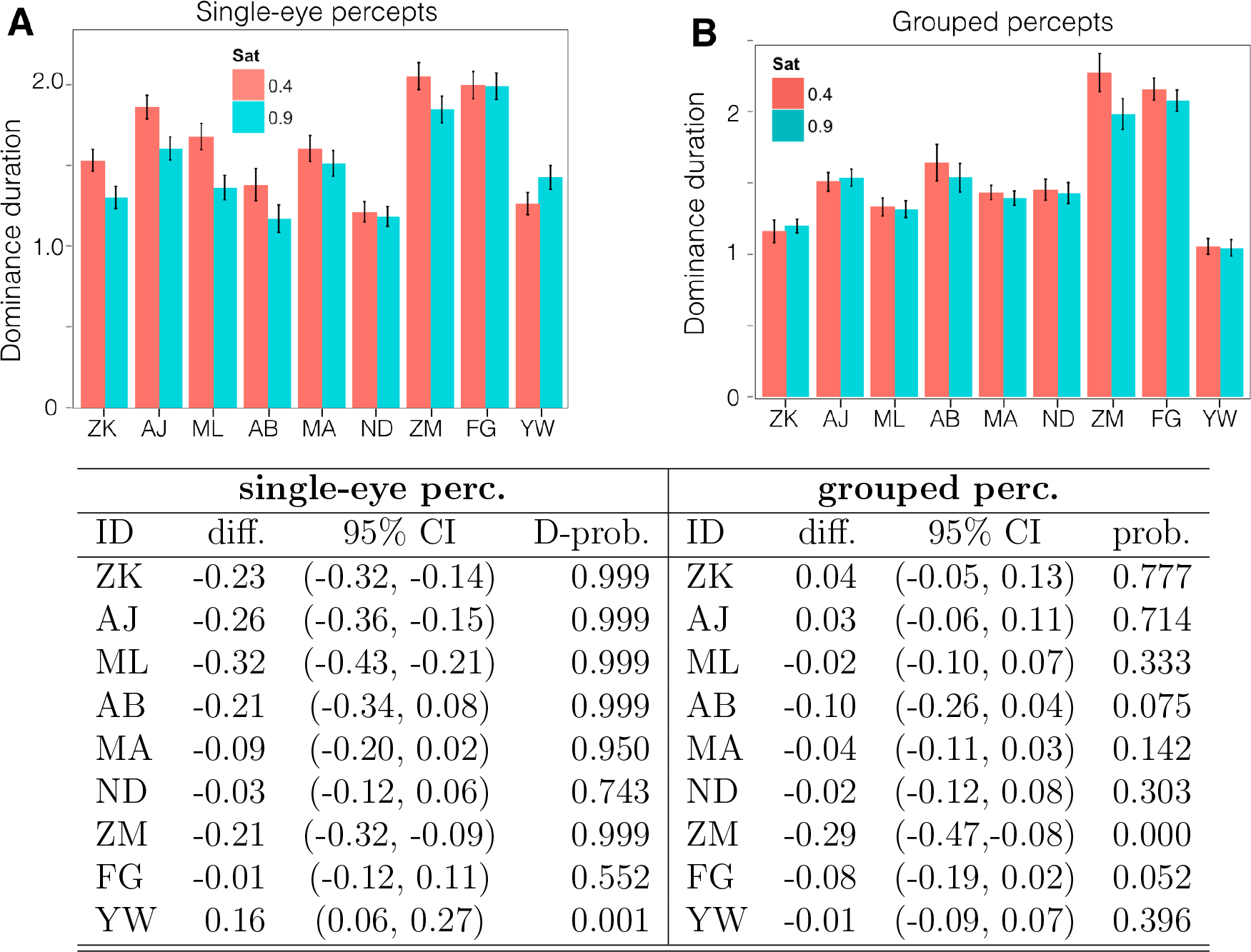
Average dominance durations: (A) single-eye percepts and (B) grouped percepts. Single-eye percept dominance durations decrease as color saturation is increased for the subjects who also experience increased grouped percept predominance. Here ‘D-prob.’ (on left) is the probability that the dominance duration of single-eye percepts decreases and ‘prob.’ (on right) is the probability that the dominance duration of grouped percepts increases.

*Grouped percept visit frequency increases with color saturation.* With multiple percepts, each can occur with a frequency between 0% to 50%, excluding self-transitions. This frequency impacts predominance (Naber et al., 2010; Huguet et al., 2014). We therefore examined how the frequency of visits to single-eye and grouped percepts depended on color saturation.

**Figure 6:**
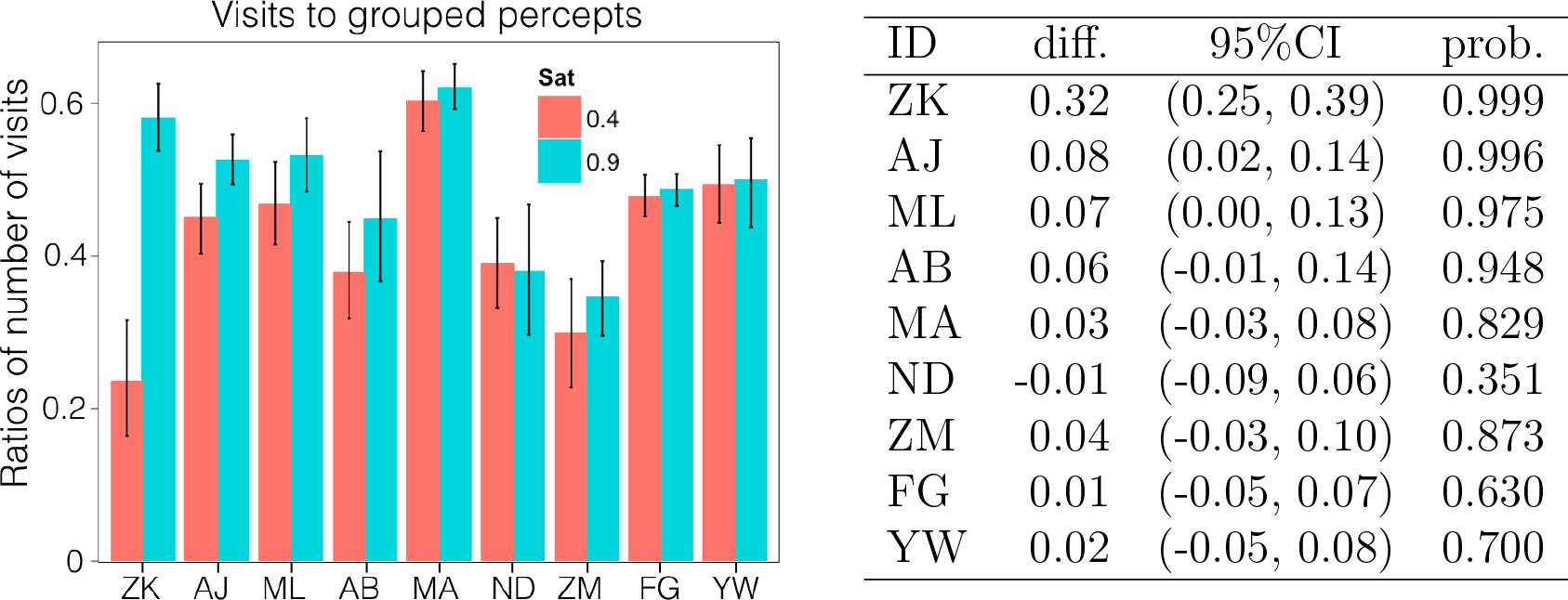
Frequency of visits to grouped percepts out of all visits. The mean increases for eight out of nine subjects when color saturation is increased from 0.4 to 0.9. The five subjects who experienced an increase in grouped percept predominance, also showed an increase in the frequency grouped percept visits. Values in the table are computed in the same way as in Fig. 4.

Consistent with our results for grouped percept predominance (Fig. 4), the frequency of visits to grouped percepts increased with color saturation in most subjects (Fig. 6, see Methods for details about the analysis): Subjects ZK, AJ, ML and AB (probability>0.94), and to a lesser degree MA (prob>0.82), show a consistent increases in the number of visits to grouped percepts.

We conclude that two main factors contributed to increased predominance of grouped percepts with increased color saturation in the five subjects affected by this change: First, the average dominance duration of single-eye percepts decreased, while the dominance durations of grouped percepts remained approximately unchanged. Second, the grouped percepts were visited more frequently in the high color saturation condition.

### 3.2 Transitions to grouped percepts increase with color saturation

We also analyzed the transition probability between percepts. We focused on the frequency of transitions between each percept type: single-eye or grouped percepts (See Fig. 7A). In doing so, we reduced the number of possible transitions to four: single-eye to grouped, grouped to single-eye, grouped to grouped, and single-eye to single-eye (See Methods). Our analysis of the frequency of visits to grouped percepts (Fig. 6) suggests an increase in transitions to grouped percepts in the high color saturation condition. Consistent with this trend, we found that there was high probability of a decrease in the ratio of transitions from single-eye to single-eye percepts for the first five subjects (ZK, AJ, ML, MA, and ZM in Fig. 7B), here ratio of 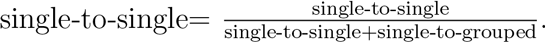. Theprobability of this decreasing effect was higher than 0.98 for four of those five subjects and higher than 0.87 for the one remaining. This implies that the ratio of the transitions from single-eye to grouped percepts increased as color saturation increased. In addition, there was high probability that the ratio of grouped percepts to grouped percepts transitions increased as the color saturation for four out of those five subjects (prob>0.94), see Fig. 7C. Thus, there was an increase in the frequency of extended bouts of grouped percepts, whereby each switch yields the opposing grouped percept. This phenomenon has previously been referred to as “trapping”, as it suggestsa subject’s perception is trapped in a subset of all possible percepts (Suzuki and Grabowecky, 2002).

**Figure 7:**
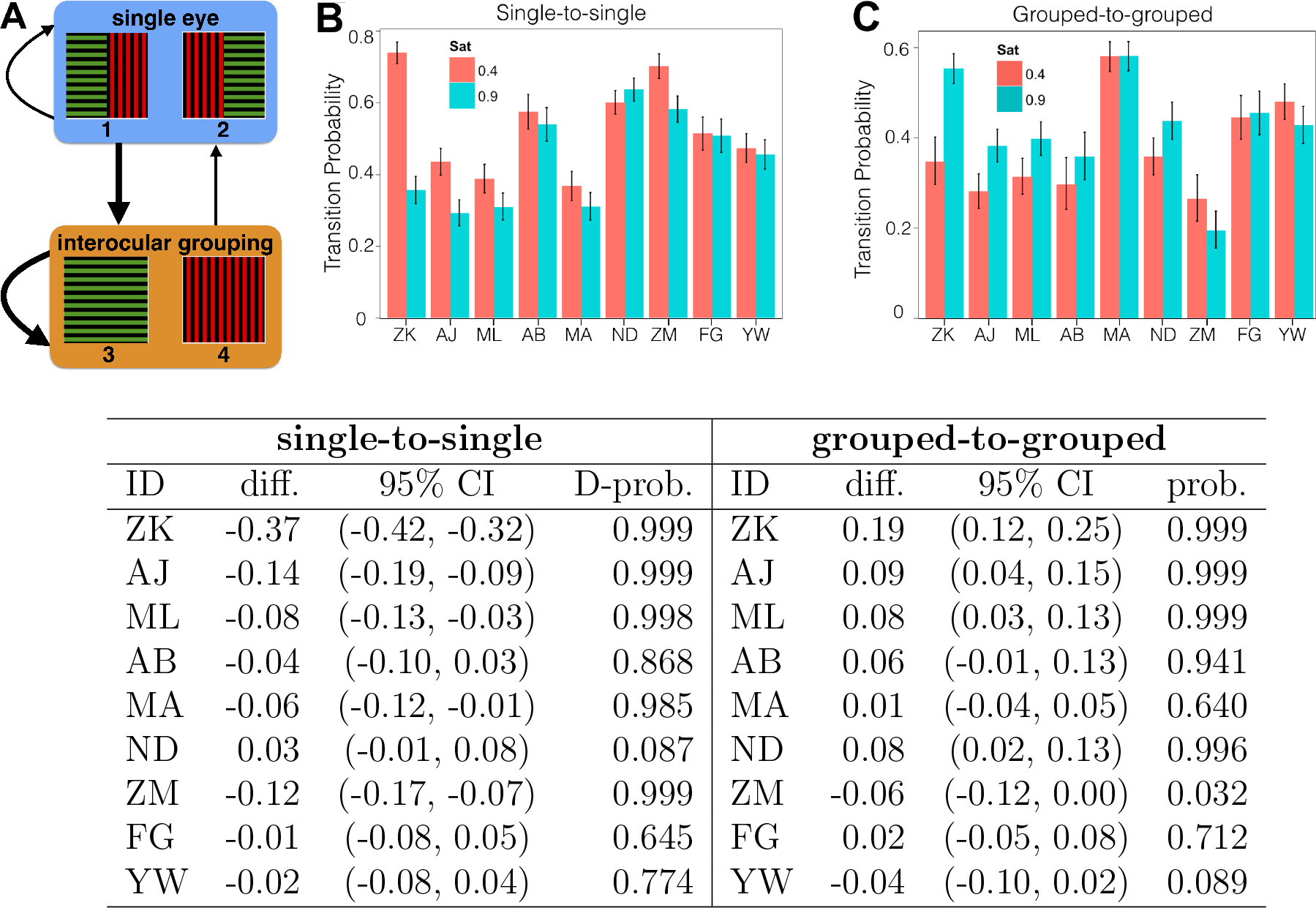
(A) Diagram showing the case where single-to-single percept transitions are less likely than grouped-to-grouped transitions, represented by the thickness of transition arrows. (B,C) The probability of transitions from (B) single-to-single percepts, and (C) grouped-to-grouped percepts.The probability of a single-to-single transition tends to decrease with color saturation whereas the grouped-to-grouped transition probability tends to increase in the cohort of subjects whose grouped predominance increased. The table gives the posterior probability of a decreases in single-to-single transition, and an increase in grouped-to-grouped transitions given the data.

### 3.3 Network mechanisms in a computational rate model

Our experimental findings suggest that increased color saturation can facilitate the binding of complementary image halves presented to either eye (as in Fig. 1). This is consistent with previous experiments demonstrating that collinear patches are grouped more frequently than orthogonal patches (Alais and Blake, 1999). Furthermore, there is evidence that collinear facilitation is influenced by chromatic cues (Huang et al., 2007).

To examine the neural mechanisms that underlie these observations, we have constructed a model of neural activity whose dynamics mirror our experimental findings (Fig. 2). The network consists of two levels, each containing four neural populations. The populations at the second level correspond to the four possible percepts. Those at the first level each correspond to the stimulus in one of the visual hemifields. Recurrent excitation between populations encoding complementary image halves is facilitated by an increase in color saturation, basedon the observation that color saturation reinforces interocular grouping (Nothdurft, 1993; Hadjikhani et al., 1998). This leads to an increased chance that populations

**Figure 8:**
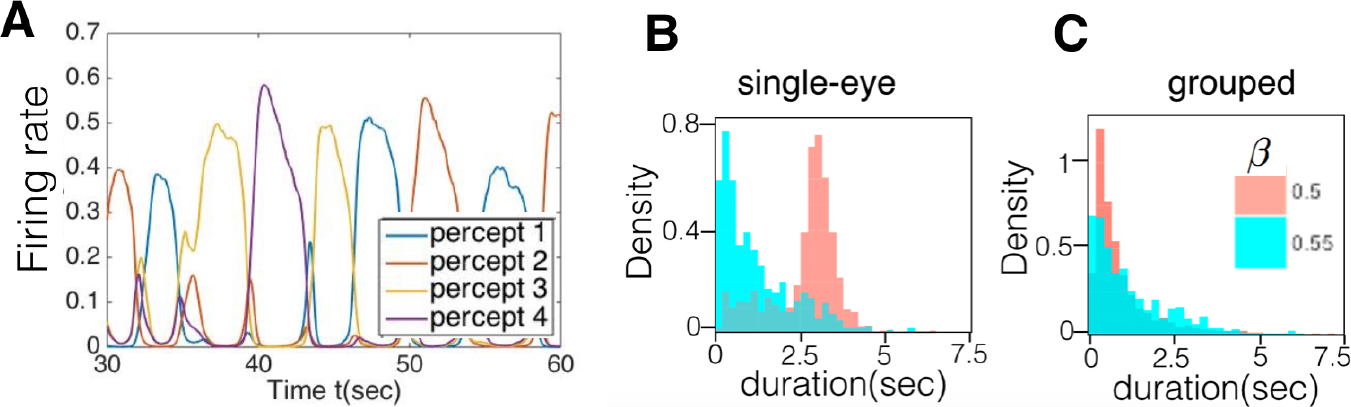
Results of a computational model of interocular grouping in perceptual multistability. (A) Numerical simulation of the model described in Fig. 2. Each trace represents the activity of a different population at Level 2, corresponding to one of four possible percepts. Note the variability in the order and timing of activations. (B,C) Dominance time distributions of (B) single-eye and (C) grouped percepts at the different values of β with fixed α=0.6 (See Supplementary Material for other parameter values).

**Figure 9:**
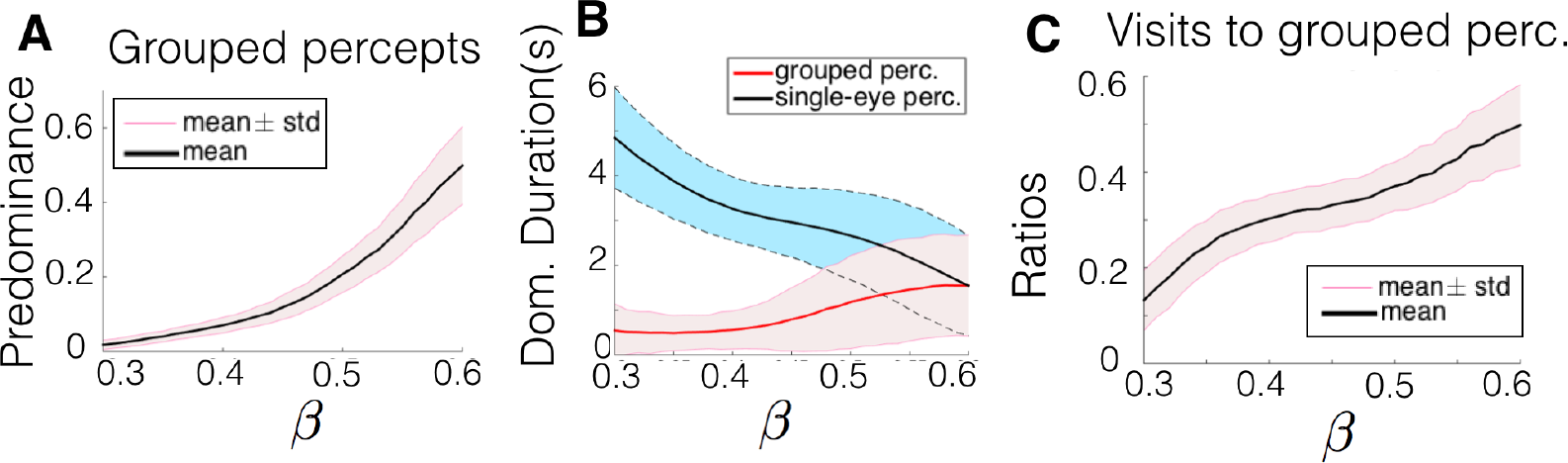
Effects of changes in color saturation, modeled by varying thecoupling parameter, α. (A) The predominance of grouped percepts increases with fi, consistent with experimental data presented in Fig. 4. (B) The average dominance duration of single-eye percepts decreased with β while that of grouped percepts remained approximately unchanged, in accordance with experimental data in Fig. 5A. (C) Furthermore, the frequency of visits to grouped percepts increased with fi, as in experimental data in Fig. 6. representing grouped images will be active.

As can be seen in a typical numerical simulation (See Fig. 8A), a single percept-related population (in Level 2) is active at any given time. Both the ordering and timing of percept dominance were stochastic, but the dominance time distributions were unimodal (Fig. 8B,C). As in previous models of perceptual multistability (Laing and Chow, 2002; Wilson, 2003; Moreno-Bote et al., 2007), we assumed perceptual switching is governed both by a slow adaptation variable as well as internal noise (See Methods). In classical bistable models of perceptual rivalry, stimulus strengths are represented by input I (or I_i_). However, in the multistable perceptual rivalry model involving interocular grouping, changing any input strength I_i_ influences the strengths of single-eye and grouped percepts simultaneously. Thus in our model, the effects of color saturation were controlled by a single parameter, β, which scaled the strength of excitatory coupling between populations responding to complementary image halves of grouped percepts. Another parameter, αrepresented the strength of excitatory coupling between populations receiving input from the hemifields of the same eye. In accordance with our experimental findings (Fig. 3), as β was increased the dominance time distributions for single-eye percepts shifted left, while the distribution of grouped dominance times remained approximately unchanged.

**Figure 10:**
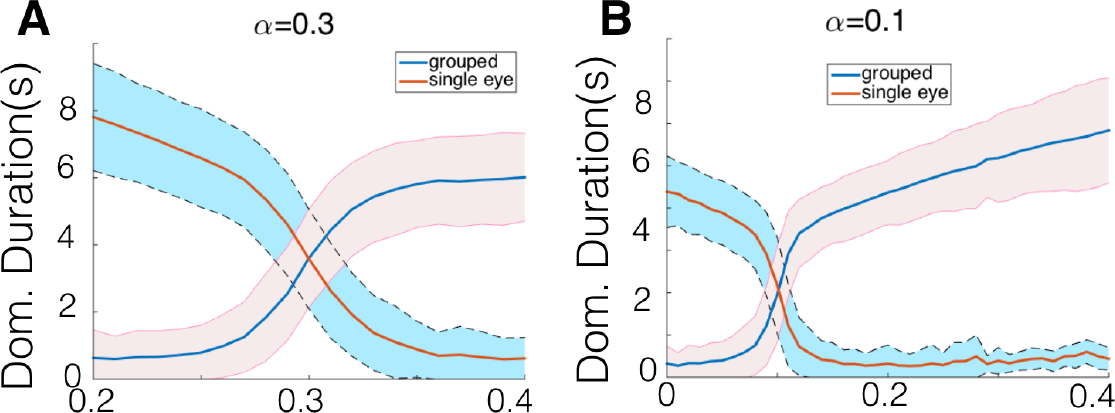
Levelt’s proposition II also holds when the coupling strength (α) between single-eye halves issmall (α=0:1) or intermediate (α=0:3).

**Figure 11:**
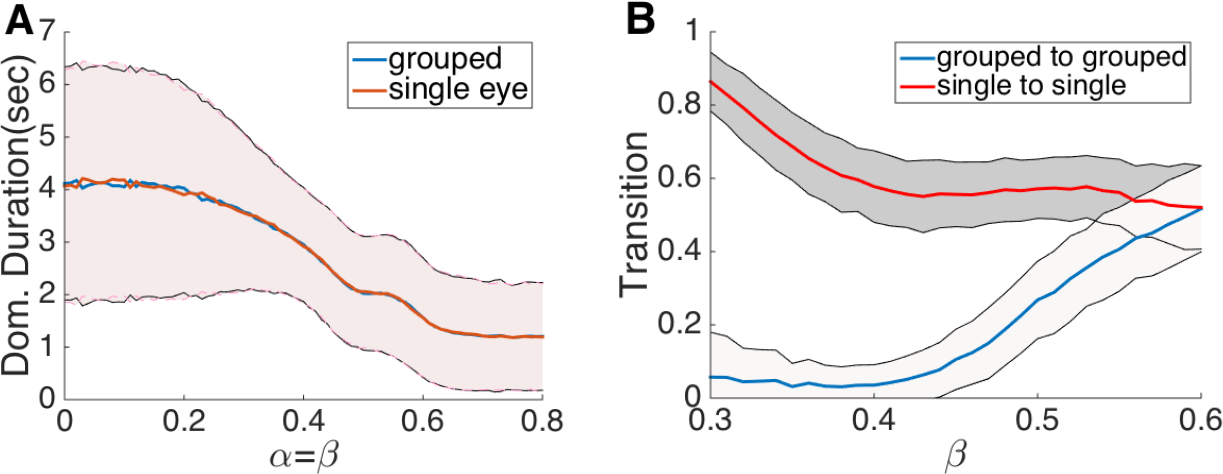
(A) Levelt Proposition IV: As the recurrent coupling between both ipsilateral and contralateral hemifield populations increases (both α and β), the dominance durations of both types of percepts (single-eye and grouped) decrease. Since these coupling strengths correspond to both percept strengths, this trend is consistent with Proposition IV. (B) The mean fractions of the transition rates (grouped to grouped and single-eye to single-eye). In simulations α=0.6 (See Methods for other parameter values).

*Single-eye percept dominance times decrease with color saturation.* Our computational network model recapitulates the three main observations of our psychophysical experiments. First, as β, the parameter representing color saturation dependent coupling, is increased the predominance of grouped percepts increased (Fig. 9A). This is consistent with the generalized Levelt’s Proposition I (see Introduction): Increasing percept strength of grouped percepts increases the predominance of those percepts. As with our experimentaldata, we explored what factors contributed to this change in predominance by computing average dominance time durations as well as the number of visits to each type of percept. When fixing α=0.6, and varying β from 0 to 0.6, the average dominance duration of single-eye percepts decreases while that of grouped percepts shows much less change and the variation is very small when β E [0.5, 0.6]. This is consistent with experimental data and the generalized Levelt’s Proposition II: Increasing the difference between the percept strength of grouped percepts and that of single-eye percepts will increase the average perceptual dominance duration of the stronger percepts. Proposition II also holds for intermediate (α=0.3 Fig. 10(A)) and small (α=0.1 Fig. 10(B)) α-values, which we were not able to test experimentally.

Thus, as in our experiments, the two main factors contributing to an increase in the predominance of grouped percepts were a reduction in single-eye percept durations, and an increase in the number of visits to grouped percepts. Therefore the network mechanisms reenforcing the collective activation of populations representing complementary image halves can explain our experimental observations.

We also note that the generalization of Proposition IV holds in our model ( See Fig. 11A). The Proposition states that the alternation rate increases as the strengths of both grouped and single-eye percepts are increased while keeping it equal. Since α and β correspond to the strength of single-eye and grouped percepts, respectively, we expect that increasing them both equally ( keeping β=α) should increase the perceptual alternation rate. This occurred over a substantial range of both parameters. Overall, our computational model suggests a combination of neural mechanisms that can account for a generalization of Levelt’s Propositions to interocular grouping.

*Transitions to grouped percepts.* We were also able to recapitulate the experimentally observed effect of color saturation on the frequency of transitions to grouped percepts. In subjects where color saturation affected interocular grouping, increasing color saturation increased the probability of perceptual transition to grouped percepts (Fig. 7). In our computational model we found that increasing β lead to both a decrease in the probability of transitions to single-eye percepts, as well as an increase in transitions to grouped percepts (Fig. 11B), consistent with our experimental observations.

## 4. Discussion

Multistable perceptual phenomena have long been used to probe the mechanisms underlying visual processing (Leopold and Logothetis, 1999). While binocular rivalry is used most frequently (Blake and Logothetis, 2002), different insights can be obtained by employing visual inputs that are integrated to produce interocularly grouped percepts (Kovacs et al., 1996; Suzuki and Grabowecky, 2002). These experiments are particularly informative when guided by Levelt’s Propositions, originally developed in the case of binocular rivalry (Levelt, 1965; Brascamp et al., 2015). Here we used this approach to identify how color saturation influences the dynamics of perceptual multistability involving interocular grouping.

*Color saturation facilitates grouping of complementary image halves.* We demonstrated that increasing the color saturation of ambiguous visual inputs can increase the predominance of grouped percepts. This is consistent with the Gestalt law of similarity (Wagemans et al., 2012) and previous work, demonstrating that color cues aid in the grouping of complementary parts of a visual object (Kim and Blake, 2004; Roelfsema, 2006). There is also evidence that the neural mechanisms underlying collinear facilitation for chromatic and achromatic contours are different (Beaudot and Mullen, 2003; Huang et al., 2007), suggesting there are multiple channels in the visual system that affect the grouping of image halves in Fig. 1. Ultimately, we propose that color provides one cue that promotes the grouping of objects between eyes.

It is important to note that we only observed an appreciable increase in grouped percept predominance in five out of nine subjects (Fig. 4). In the remaining four subjects we did not observe an effect of color saturation on percept predominance. One possible reason for this result is that subjects differed in their sensitivity to color saturation (Kaiser and Boynton, 1996). Although no subjects reported problems with distinguishing colors, they may have responded differently if the change in color saturation was larger, or if we used different colors. For example, the wide array of sensitivities to contrast across human subjects are reflected in the range of mean dominance time durations in binocular rivalry (Bossink et al., 1993; Brascamp et al., 2006; van Ee, 2009). Also, the relationship between color saturation and percept predominance is likely nonlinear Stalmeier and de Weert (1998). The color saturation values we used may have fallen in the flat portion of the function that describes the relation between color saturation and predominance for the four unaffected subjects.

As mentioned previously, Stalmeier and de Weert found significant inter-subject variability even when isoluminance points were calibrated individually for each subject Stalmeier and de Weert (1998). The effect of chromatic signal strength on binocular rivalry depended both on the calibration criterion (flicker photometry versus MDB) and the direction along which colors are sampled in the color space. Stalmeier and de Weert also showed significant inter-subject variability both in the absolute effectiveness of achromatic contrast and its relative effectiveness with respect to chromatic contrast Stalmeier and de Weert (1998). Inter-subject variability has been reported in relatively low-level tasks (e.g. (Halpern et al., 1999)), as well as in multistable perception (Kleinschmidt et al., 2012), which is interpreted to include both low-level and high-level factors. Hence, for future studies, we suggest the use of multiple levels of the percept-strength variable in order to characterize more completely the performance of each subject individually. In addition, it would help us identify the relative contributions of color saturation and luminance to percept strength, since red and green have different luminance at a fixed saturation (See Methods). This would provide a test for the generality of our conclusions. Increasing the number of subjects would allow us to better characterize inter-subject variability, but would likely not make it disappear.

*Extending Levelt’s propositions to interocular grouping.* Interocular grouping has been reported with different sets of patchwork images (Kovacs et al., 1996; Suzuki and Grabowecky, 2002). However, earlier studies did not quantify specific ways in which a stimulus parameter could affect the predominance of grouped images. We have shown that color saturation used as a grouping cue differentially controls the strength of single-eye and grouped percepts, and increasing color saturation can increase grouped percept predominance. This suggests color saturation may act as a stimulus strength parameter for grouped percepts, in line with Proposition I.

In agreement with Proposition II, the single-eye percepts began with higher predominance, and their dominance durations were decreased in the higher color saturation condition. Consistent with Proposition III, we found the average dominance duration decreased. Finally, since we could not determine whether we equally increased the strength of both single-eye and grouped percepts, it is unclear whether our results are consistent with Levelt Proposition IV. Color saturation may affect monocular and binocular integration in different ways (Sincich and Horton, 2005). Stimulus parameter changes obeying Proposition IV would have to keep predominance fixed, while decreasing mean dominance durations. However, in our computational model, we find that when single-eye and interocular coupling strengths are equal,*i.e.* α=β, then increasing both leads to a decrease in dominance time durations (See Fig. 11A).

Studies of interocular grouping in perceptual multistability have a long history (Diaz-Caneja, 1928). We focused on split single-eye images for simplicity, but we anticipate that our findings extend to the patchwork images of Kovacs et al. (1996). The simple grating-based inputs we used were more similar to the geometric images of Suzuki and Grabowecky (2002). We expect that our findings extend to achromatic images as long as a parameter can be identified that affects grouped percept predominance. For example, we could use achromatic textures as a cue to group complementary stimulus halves. In general, we suggest that our findings apply to any stimulus feature that promotes grouping along the lines of Gestalt laws of grouping.

*Extensions to other computational models.* We made several specific choices in our computational model. First, we described neural responses to input in each visual hemifield by a single variable. We could also have partitioned population activity based on orientation selectivity or receptive field location (Ferster and Miller, 2000). This would allow us to describe the effects of horizontal connections that facilitate the representation of collinear orientation segments in more detail (Bosking et al., 1997; Angelucci et al., 2002). Since there is evidence for chromatically-dependent collinear facilitation (Beaudot and Mullen, 2003), we could model the effects of image contrast and color saturation as separate contributions to interocular grouping. In the present model, the effects of color saturation were described by a single parameter, β, representing the coupling between the neural populations at the first level of our hierarchy. Increasing color saturation could have also impacted the stimulus strength I, e.g., through changes in luminance. A more detailed study of the effects of increasing both β and I to account for saturation and luminance changes will be pursued in future work. We expect that color saturation also affects neural activity and contextual feedback in higher visual areas (Sincich and Horton, 2005). We could therefore extend our model to account for chromatic effects on the activity at the second level in Fig. 2.

*Comparisons with previous models of perceptual multistability.* Our computational model is based on the assumption that perceptual multistability occurs via a winner-take-all process, with a single percept temporarily excluding all others (Wilson, 2003; Shpiro et al., 2007). Subsequently, some neural process must allow the system to switch from the dominant to another percepts after a few seconds (Laing and Chow, 2002). The simplest mathematical model of this process is a multistable system where slow adaptation and/or noise drives switches between multiple attractors (Moreno-Bote et al., 2007; Braun and Mattia, 2010). This framework is common in models of binocular rivalry (Laing and Chow, 2002; Shpiro et al., 2007), non-eye-based perceptual rivalry (Brascamp et al., 2009), and even perceptual multistability with more than two percepts (Diekman et al., 2013; Kilpatrick, 2013; Huguet et al., 2014). Each percept typically corresponds to a single neural population which mutually inhibits the other(s). Spike rate adaptation or short term plasticity then drive the slow switchingbetween networkattractors (Laing and Chow, 2002), and noise generates variation in the dominance times (Moreno-Bote et al., 2007).

Our computational model differs from previous ones in a few key ways. First, excitatory connectivity at the first level facilitates both single-eye and grouped binocular percepts. Second, there is a hierarchy of levels,with percepts represented by neural activity at the higher level. Importantly, the strength of excitatory connectivity at the first level determines the input strength to second level populations, and ultimately each percept’s predominance. In this way, our model is similar to that of Wilson Wilson (2003), who used a two level model to capture the effects of monocular and binocular neurons. However, Wilson’s model focused on the case of two possible percepts, while our computational model accounts for all four possible percepts in an interocular grouping task.

*Neural mechanisms of perceptual multistability.* Our observations support the prevailing theory that perceptual multistability is significantly percept-based and involves higher visual and object-recognition areas (Leopold and Logothetis, 1999). Since the first systematic study on binocular rivalry (Wheatstone, 1838), much work has been devoted to identifying its underlying neural mechanisms: Mutual inhibition allows for the selection of one percept among many (Lumer, 1998; Tong and et al, 1998; Tong, 2001; Lee et al., 2005; Haynes et al., 2005; Meng et al., 2005; Moutoussis et al., 2005; Wunderlich et al., 2005; Seely and Chow, 2011), adaptation can lead to switching between percepts (Kim et al., 2006; Brascamp et al., 2006; van Ee, 2009), and neuronal noise accounts for the irregularity of perceptual dominance intervals (Brascamp et al., 2006; Moreno-Bote et al., 2007; Shpiro et al., 2009; Lankheet, 2006). However, a number of issues remain unresolved. Activity predictive of a subject’s dominant percept has been recorded in lateral geniculate nucleus (LGN) (Haynes and Rees, 2005), primary visual cortex (V1) (Lee and Blake, 2002; Polonsky et al., 2000), and higher visual areas (e.g., V2, V4, MT, IT) (Logothetis and Schall, 1989; Leopold and Logothetis, 1996; Sheinberg and Logothetis, 1997). Thus, rivalry likely results from interactions between networks at several levels of the visual system (Freeman, 2005; Wilson, 2003). As a result, it is important to develop descriptive models that incorporate multiple levels of the visual processing hierarchy.

Collinear facilitation involves both recurrent connectivity in V1 as well as feedback connections from higher visual areas like V2 (Angelucci et al., 2002; Gilbert and Sigman, 2007), reenforcing the notion that perceptual rivalry engages a distributed neural architecture. However, a coherent theory that relates image features to dominance statistics during perceptual switching is lacking. It is unclear how neurons that are associated to each subpopulation may interact due to grouping factors such as collinearity and color.

Although we only presented one model in this work, we have tested two others: one model assumes that interocular grouping is encoded by strengthening the feedforward coupling from the Level 1 to Level 2, and the another model assumes that grouping effects occur via stronger feedback from Level 2 to Level 1. Interestingly, all three models produce similar results.

*Conclusion.* Our work supports the general notion that perceptual multistability is a distributed process that engages several layers of the visual system. Interocular grouping requires integration in higher visual areas (Leopold and Logothetis, 1996), but orientation processing and competition occurs earlier in the visual stream (Angelucci et al., 2002; Gilbert and Sigman, 2007). Furthermore, the fact that color saturation can modulate the statistics of perceptual multistability provides a novel stimulus parameter that can be varied in visual inputs to probe the neural mechanisms of visual integration and competition.

## 5. Acknowledgments

Gemma Huguet and Ruben Moreno-Bote provided helpful comments. Funding was provided by NSF-DMS-1311755 (ZPK); NSF-DMS-1517629 (KJ and ZPK); and NSF-DMS-1122094 (KJ). YW was supported by DHS-2014-ST-062-000057 and by a seed grant from Texas Southern University.

Supplementary material can be found at https://github.com/YunjiaoWang/multistableRivalry.git

